# Discovery of a Promising Susceptibility Factor for Fusarium Head Blight in Wheat

**DOI:** 10.1101/2020.11.19.390534

**Authors:** Bhavit Chhabra, Vijay Tiwari, Bikram S Gill, Yanhong Dong, Nidhi Rawat

## Abstract

Fusarium head blight (FHB) disease of wheat caused by *Fusarium* spp. deteriorates both quantity and quality of the crop. Manipulation of susceptibility factors, the genes facilitating disease development in plants, offers a novel and alternative strategy for enhancing FHB resistance in plants. In this study, a major effect susceptibility gene for FHB was identified on the short arm of chromosome 7A (7AS). Nullisomic-tetrasomic lines for homoeologous group-7 of wheat revealed dosage effect of the gene, with tetrasomic 7A being more susceptible than control Chinese Spring wheat, qualifying it as a bonafide susceptibility factor. The gene locus was conserved in six chromosome 7A inter-varietal wheat substitution lines of diverse origin and a tetraploid *Triticum dicoccoides* genotype. The susceptibility gene was named as *SF^7AS^FHB* and mapped on chromosome 7AS to 48.5-50.5 Mb peri-centromeric region between del7AS-3 and del7AS-8. Our results showed that deletion of *SF^7AS^FHB* imparts ~ 50-60% type 2 FHB resistance (against the spread of the fungal pathogen) and its manipulation may lead to enhanced resistance against FHB in wheat.

**Highlight:** Discovery and mapping of a conserved susceptibility factor located on the short arm of wheat chromosome 7A whose deletion makes plants resistant to Fusarium Head Blight.

## Introduction

Fusarium Head Blight (FHB) is one of the most destructive diseases of wheat worldwide affecting the yield and safety of grain. Severe outbreaks of FHB may cause up to 50% yield loss, in addition to the serious risk to grain quality because of the associated mycotoxins (Snijders, 1990, Parry et al., 1995). In the USA alone, during 1991-1996, FHB outbreaks caused an estimated $3 billion crop loss (McMullen et al., 1997) During the 2015-16 crop year, economic loss due to FHB was estimated at $1.2 billion (Wilson et al., 2018).

FHB is caused by a complex of *Fusarium* species (Parry et al., 1995). In North America, the predominant causal organism is *Fusarium graminearum* (McMullen et al., 1997; Goswami & Kistler, 2004). In wheat, typical symptoms are pre-mature bleaching of infected spikelets resulting in aborted or shriveled seeds and hence, reduced yield. Associated mycotoxins, such as deoxynivalenol (DON) severely reduce quality of the grain (Snijders, 1990; Chen et al., 2019). DON is phytotoxic and can cause wilting, chlorosis and necrosis (Cutler, 1988) as well as inhibits protein synthesis in mammals by binding to 60S subunit of eukaryotic ribosomes (Rocha et al., 2005). US Food and Drug Administration (FDA) has set an advisory limit of 1 ppm DON for wheat and barley products for human consumption, 10 ppm for cattle and poultry, 5ppm for swine and all other animals (https://www.fda.gov/regulatory-information/search-fda-guidance-documents/guidance-industry-and-fda-advisory-levels-deoxynivalenol-don-finished-wheat-products-human accessed Oct 5, 2020). In wheat, DON acts as a virulence factor for *Fusarium*, helping the pathogen in disease spread (Bai et al., 2002; Jansen et al., 2005).

Integrated management practices incorporating genetic resistance, chemical and agronomic control measures are used for controlling FHB (Wegulo et al., 2010; Salgado et al., 2014). Genetic resistance is the most economical, environment-friendly and effective component of the overall strategy to control FHB (Bai & Shaner, 2004). Mesterházy et al. (1999) described five types of genetic resistance to FHB, out of which, Type 1 (resistance to initial infection) and Type 2 (resistance to spread within the spike) are the most widely studied in wheat. Type 2 resistance is less affected by non-genetic variables as compared to Type 1 resistance (Bai & Shaner, 1994). Resistance against FHB is quantitively controlled. *Fhb1*, the first Quantitative Trait Locus (QTL) for resistance against FHB in wheat was discovered in 1999 (Waldron et al., 1999). Since then, around 500 QTL (104 major effect) have been reported in the literature (Buerstmayr et al., 2019). Recently another FHB resistance QTL, *Fhb7*, was characterized (Wang et al., 2020). Even after decades of efforts, achieving high levels of FHB resistance in wheat varieties is a challenge.

Polyploid nature of wheat offers it with a unique feature of genome buffering, due to which wheat can tolerate addition, substitution or deletion of different sets of chromosomes and still be fertile (Sears, 1954; Endo, 2015). A number of aneuploid stocks have been developed in bread wheat by wheat cytogeneticists using this feature that have served as a very important genetic resource for the wheat community (Sears, 1954; Sears, 1966; Endo and Gill, 1996; for review see Gupta & Vasistha, 2018). Different types of aneuploid stocks developed by ER Sears over the years were: Monosomics (one chromosome lost), Trisomics (one chromosome gained), Nullisomics (one pair of homologous chromosomes lost), Tetrasomics (one pair of homologous chromosomes gained), Nullisomic-tetrasomics (tetrasome of a homeolog group compensates for the nullisome for each of the other two homelog chromosomes of same group) and Di-telosomics (lack chromosome arms for one homologous group) (Sears, 1954; Sears, 1966). Endo & Gill (1996) reported isolation of 436 Deletion lines across all chromosomes of Chinese Spring with random break points by utilizing an alien chromosome from *Aegilops* spp. All these genetic stocks have been developed in Chinese Spring wheat background and have led to its establishment as the reference wheat variety, in spite of its poor agronomic traits. Cytogenetics-based physical maps for all seven homoeologous groups were developed using these stocks (Werner et al. 1992; Endo 1986; Endo and Gill 1996). All seven of the Mendelized QTL reported for FHB have used these cytogenetic stocks to find either candidate chromosome, chromosome arm or to refine the candidate gene region (Cuthbert et al., 2006, 2007; Qi et al., 2008; Xue et al., 2010, 2011; Cainong et al., 2015; Guo et al., 2015). Ma et al. (2006) screened a set of 30 ditelosomic (Dt) lines of Chinese Spring for their response to *F. graminearum* infection and found that they varied in their response to FHB. Five out of thirty lines (Dt 6BS, Dt 4DL, Dt 7BL, Dt 3BS and Dt 7AL) had lower proportion of scabby spikelets (p<0.01) and four lines (Dt 6AL, Dt 6DS, Dt 4DL and Dt 7AL) had significantly less DON content (p<0.05) as compared to CHINESE SPRING. The reason might be the presence of susceptibility genes or resistance suppressors present on their missing chromosome arms. Dt 7AL (missing 7AS) showed lowest amount of FHB severity as well as minimal DON content, and chromosome 7A was therefore selected in the present work for further investigation.

Plants activate phytohormone-regulated defense responses for protection against pathogens. On the other hand, pathogens release effector molecules to manipulate phytohormones and defense responses for their own benefit (Kazan & Lyons, 2014; Han & Kahmann, 2019). Effectors released by pathogens either suppress gene targets having a role in defense or activate targets that may support its growth or have negative impact on defense pathways (Pavan et al., 2009). Eckardt (2002) first introduced the term plant susceptibility factors (SF). Genetically, they can be considered as dominant genes whose manipulation will lead to recessive resistance (Pavan et al., 2009). In common language, genes facilitating infection or supporting compatibility of plant-pathogen interaction are termed as susceptibility genes. Hence mutation in them limits the pathogen spread. Susceptibility genes can be divided into three categories based on their role during different stages of infection: a)- Helping pathogen in establishment; b)- Involved in modulating/regulating plant defenses c)- Involved in providing nutrition to the pathogen (van Schie & Takken, 2014).

Manipulating susceptibility genes by knocking out or knocking down their expression provides a novel and alternative breeding strategy for protection against pathogens. Benefits that come along with utilizing susceptibility genes are that the resistance acquired is recessive, broad spectrum and more durable as compared to R-genes. However, as most of the susceptibility genes have pleiotropic effect hence manipulating them may affect the host’s physiological processes. A well-known example of a susceptibility gene used in crop breeding is Barley *mlo* gene, this loss of function mutation has provided broad spectrum resistance against powdery mildew for over 35 years in many plant species (Büschges et al., 1997; Engelhardt et al., 2018).

In the present work, we identified a major susceptibility factor for FHB on chromosome 7A of wheat, and further narrowed down its location to a 48.5-50.5 Mb physical interval on the short arm of chromosome 7A.

## Materials and Methods

### Plant Material

Experiments were conducted over three seasons (2018, 2019 and 2020) in Research Greenhouse Complex, University of Maryland, College Park. Plant material used in the experiments is listed in Table 1. All of the material used was in Chinese Spring genetic background, therefore, wild type Chinese Spring was used as a positive control in all the experiments. Temperature conditions were 23-25 °C during daytime and 16-18 °C during night-time. Photoperiod profile of 16hr (day): 8hr (night) was used. Five plants for each line were grown in 6” pots. Nulli-tetrasomic lines and ditelosomic lines were tested only in year 2020 whereas deletion and substitution lines were analyzed in all three sets (2018, 2019, 2020). Del7AS-6 and Del7AS-3 could not be tested in year 2019 because of technical mishaps. A subset of deletion lines critical for locating *SF^7AS^FHB* was tested again in an additional set in 2020 along with control Chinese spring.

**Table 1:** Description of plant material used in the study

| Accession<sup>a</sup> | Description | Abbreviation<sup>b</sup> | Source |
| --- | --- | --- | --- |
| TA 3282 | Chinese Spring Nulli 7A- Tetra 7B | CS N7A-T7B | WGRC |
| TA 3283 | Chinese Spring Nulli 7B- Tetra 7A | CS N7B-T7A | WGRC |
| TA 3285 | Chinese Spring Nulli 7D- Tetra 7A | CS N7D-T7A | WGRC |
| TA 3110 | Chinese Spring Ditelosomic 7AS | CS Dt7AS | WGRC |
| TA 3111 | Chinese Spring Ditelosomic 7AL | CS Dt7AL | WGRC |
| TA 3121 | Chinese Spring Ditelosomic 7BS | CS Dt7BS | WGRC |
| TA 3122 | Chinese Spring Ditelosomic 7BL | CS Dt7BL | WGRC |
| TA 3130 | Chinese Spring Ditelosomic 7DS | CS Dt7DS | WGRC |
| TA 3071 | Chinese Spring Ditelosomic 7DL | CS Dt7DL | WGRC |
| TA 3221 | Chinese Spring-Timstein Disomic Substitution 7A T(7A CS) | CS-T DS7A | WGRC |
| TA 3447 | Chinese Spring-Dicoccoides Disomic Substitution 7A TDIC(7A CS) | CS-TDIC DS7A | WGRC |
| TA 3242 | Chinese Spring-Cheyenne Disomic Substitution 7A CNN(7A CS) | CS-CNN DS7A | WGRC |
| TA 3782 | Chinese Spring-Thatcher Disomic Substitution 7A TH(7A CS) | CS-TH DS7A | WGRC |
| TA 3200 | Chinese Spring-Hope Disomic Substitution 7A H(7A CS) | CS-H DS7A | WGRC |
| TA 3451 | Chinese Spring-Axminster Disomic Substitution 7A AM(7A CS) | CS-AM DS7A | WGRC |
| TA 4536 | Chinese Spring Deletion 7AS-1 | CS del7AS-1 | WGRC |
| TA 4546 | Chinese Spring Deletion 7AS-2 | CS del7AS-2 | WGRC |
| TA 4546 | Chinese Spring Deletion 7AS-3 | CS del7AS-3 | WGRC |
| TA 4546 | Chinese Spring Deletion 7AS-4 | CS del7AS-4 | WGRC |
| TA 4546 | Chinese Spring Deletion 7AS-5 | CS del7AS-5 | WGRC |
| TA 4546 | Chinese Spring Deletion 7AS-6 | CS del7AS-6 | WGRC |
| TA 4546 | Chinese Spring Deletion 7AS-8 | CS del7AS-8 | WGRC |
| TA 4546 | Chinese Spring Deletion 7AS-9 | CS del7AS-9 | WGRC |
| TA 4546 | Chinese Spring Deletion 7AS-10 | CS del7AS-10 | WGRC |
| TA 4518 | Chinese Spring Deletion 7AS-11 | CS del7AS-11 | WGRC |
| TA 4546 | Chinese Spring Deletion 7AS-12 | CS del7AS-12 | WGRC |

<sup>a</sup>Accession numbers of the Wheat Genetics Resource Center (WGRC), Kansas State University (KSU).
<sup>b</sup>Abbreviations according to Raupp et al. (1995)

### Marker development, PCR conditions and Gel electrophoresis

For monitoring the identity and size of deletions in the set of deletion lines used, genome specific markers were developed every 10 Mb of 7AS arm, selecting gene sequences at 10Mb interval on the Chinese spring wheat reference genome sequence version 1.0 (IWGSC et al., 2018). To cover the entire ~360 Mb long chromosome 7AS (IWGSC et al., 2018), a total of 36 markers were designed starting from the telomeric end of the chromosome (Table 2). Genome Specific Primer (GSP) design software (Wang et al., 2016) with default settings was used for designing primers specific to chromosome 7A. A touch down Polymerase Chain reaction (PCR) profile from 64 °C to 58 °C (95 °C for 5 min, 7 cycles of 95 °C for 45 sec, 64-58 °C for 45 sec with a decrease of 1 °C per cycle, 72 °C for 1 min, followed by 27 cycles of 95 °C for 45 sec, 58 °C for 45 sec, 72 °C for 1 min, and a final extension of 72 °C for 7 min) was used. PCR product obtained was run on Ethidium Bromide stained 1.5-2% Agarose gel for 1-1.5 hr. and visualized under UV.

**Table 2:**
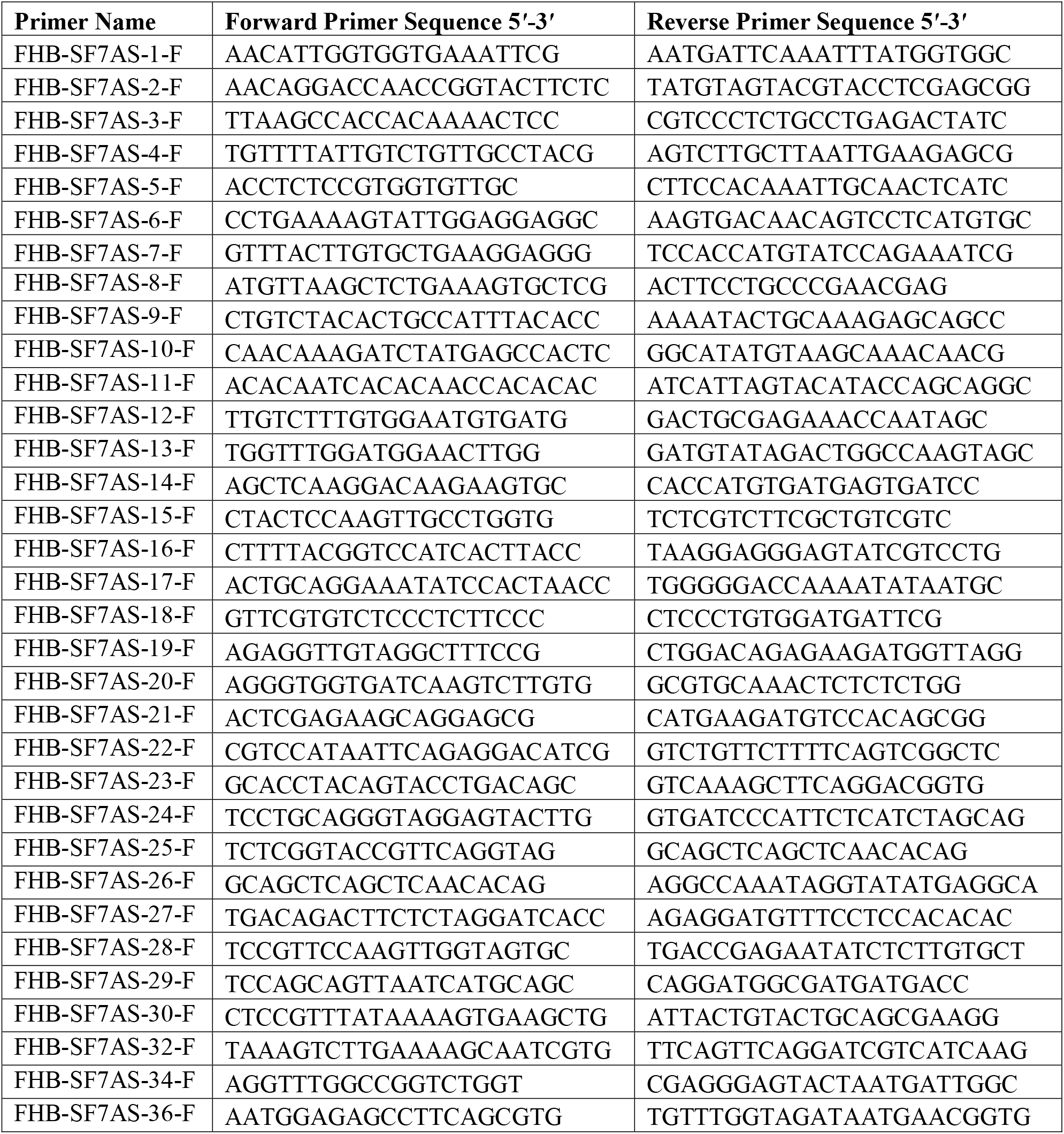
Names and sequences of genome specific primers used to genotype 7AS deletion lines

### Fungal inoculum preparation

*F. graminearum* isolate GZ3639, which is known for its strong virulence, was used in all the experiments (Desjardins et al., 1997; Rawat et al., 2016). Fungal plugs from 50% glycerol stocks (stored at −80 °C) were used to culture the fungal mycelia on Potato Dextrose Agar (PDA). Each PDA plate was inoculated with one plug from the stock, taped with autoclaved parafilm and fungus was allowed to grow at room temperature. Plates were checked for bacterial contamination each day and contaminated cultures were discarded. After seven days of inoculation, two plugs from PDA plates with actively growing fungus were inoculated on liquid Mung bean (MB) broth, shaken at 180 rpm at room temperature for a week. Macroconidia at a concentration of 1×10^5^ diluted in distilled water were used for inoculations.

### Inoculation strategy and phenotyping

Tenth and eleventh spikelets counted from the base of the spikes were point-inoculated with 10 μl macroconidial inoculum at pre-anthesis stage. Five to six spikes per plant were inoculated and covered with moisture saturated Ziplock bags for 72 hours to provide high humidity for fungal growth. Disease scoring was done 14, 21 and 28 days after inoculation (dai). Phenotyping was done by counting the number of bleached spikelets, including the inoculated spikelet, downwards from the point of inoculation

### FHB Severity, AUDPC and FDKs

Percentage of symptomatic spikelets (PSS) were calculated by converting number of bleached spikelets (the inoculation point (10th spikelet) and bleached ones down from the point of inoculation) to a scale of 100. To compare the FHB severity among different genotypes, area under the disease progress curve (AUDPC) values were calculated from the average of PSS at 14, 21 and 28 dai for each genotype. Formulae given by Simko & Piepho (2011) was used for calculating AUDPC values:

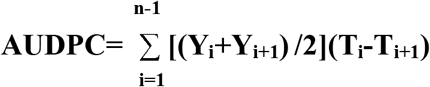

Where Y_i_ is the average of PSS (in percentage) at the ith observation, T_i_ is time (days) at the ith observation and n is the total number of observations.

Seeds were manually threshed individually from each inoculated spike after maturity and bulked by genotypes. Fusarium damaged kernels (FDKs) were measured for each genetic line by counting visually scabby kernels among all the threshed seeds of that line in the particular experiment and converting into percentage values.

### DON content

DON content of seeds was measured at USWBSI DON-testing laboratory at the University of Minnesota by GC/MS following Mirocha et al. (1998). Each sample was analyzed in three replications. Briefly, 1 g of seeds were extracted with 10 mL of acetonitrile/water (84/16, v/v) in 50 mL centrifuge tubes. Each sample was placed on a shaker for 24 hrs., and then 4 ml of the extract was passed through a column packed with C18 and aluminum oxide (1/3, w/w). Two milliliter of the filtrate was evaporated to dryness under nitrogen at room temperature, and 100 μl of Trimethylsilyl (TMS) reagent (TMSI/TMCS, 100/1,v/v) was added to the vial, rotating the vial so that the reagent makes contact with residue on the sides of the vial. The vial was placed on a shaker for 10 min, and then 1mL of isooctane containing 500ng/mL mirex was added and shaken gently. HPLC water (1ml) was added to quench the reaction and the vial was vortexed so that the milky isooctane layer became transparent. The upper layer was transferred into a GC vial for GC/MS analysis (Shimadzu GCMS-QP2020, Shimadzu Corporation, Kyoto, Japan) and readings were recorded. DON content was measured only for year 2020.

### Statistical analysis

Data was analyzed in R (vR x64 3.6.3), R studio and Excel for all sets of experiments. All the experiments were conducted in a Completely Randomized Design (CRD). Parameters analyzed were: FHB Severity, DON content, AUDPC, and FDKs. Each spike tested was considered as individual replicate. Each data set was first tested for normality and homogeneity of error variances before doing analysis. Normality was checked by plotting QQ-plots and performing Shapiro-Wilk tests. Homogeneity of error variances was checked by plotting residual vs fitted plots and performing Levene test. Experiments on nulli-tetrasomic and di-telosomic lines was performed only once in 2020, hence there is no variable year. PSS was taken as dependent variable whereas genotype was considered as independent variable with fixed effect. Kruskal-Wallis rank sum test was performed for 21 and 28 dai as dataset did not meet ANOVA assumptions of normality and homogeneity of error variances.

For FHB severity data of deletion lines and substitution lines, PSS was taken as dependent variable whereas genotype and year were considered as independent variables. Both genotype and year were considered as fixed effect. A two-way Analysis of Variance (ANOVA) with interaction was performed for 28 dai over three years. For deletion lines, Type-1 two-way ANOVA was performed as we lacked two genotypes (7AS-3 and 7AS-6) in 2019 set due to technical reasons, however, for substitution lines, type-3 two-way ANOVA was performed. Data points where significant Genotype*Year interaction was found, a simple one-way ANOVA was performed separately for each year. Type-3 one-way ANOVA was performed for both deletion and substitution lines. Data from which residuals were not normally distributed or did not appear independent of fitted values were log10 transformed, in order to meet the assumptions of data analysis.

In order to make pair-wise comparisons between control Chinese spring and other genotypes tested in each season for all the experiments, a post-hoc test (Dunnett test if analysis conducted by ANOVA or Dunn test if analysis conducted by Kruskal-Wallis rank sum test) was performed. For analysis of AUDPC values and DON content, same set of procedures was followed as that for FHB severity data of deletion lines for all the experiments. Data for DON content and AUDPC values which did not meet assumptions of ANOVA were square root transformed, in order to meet the assumptions of data analysis.

For analysis of FDK values, a single sample t-test with unknown standard deviation was used as only one replicate/ genotype was available for all experiments. First, standard deviation was calculated by considering whole data set. A critical t-value was obtained from the t-table on the basis of degrees of freedom and at α=0.05. T-value was calculated between control and other genotypes separately for each experiment.

## Results

### FHB susceptibility factor is located on chromosome 7A and has a dosage effect

It would be appropriate to call the underlying locus a susceptibility factor, if increasing the copies of 7A makes the plants more susceptible as compared to parental Chinese spring (dosage effect). For this purpose, Nullisomic-tetrasomic lines for chromosome 7A were evaluated for their response to *F. graminearum* infection.

FHB severity in the inoculated plants was recorded at 14, 21 and 28 dai, and was found to be maximum at 28 dai in Chinese spring control. Therefore, final comparisons of FHB severity were done at 28 dai in all the experiments. FHB severity of nulli-tetrasomic stocks for chromosome 7A ranged from 10 to 100% (Kruskal-Wallis test explained statistical significance at p= 7.093e^−07^ with chi-square value of 31.373. Dunn test results showed all three of tested Nulli-tetra lines to be significantly different from control. N7A-T7B showed significantly lower disease at p<0.01 whereas N7B-T7A and N7D-T7A had significantly higher disease severity at p<0.001 (Fig. 1a; Fig. 1e).

**Figure 1:**
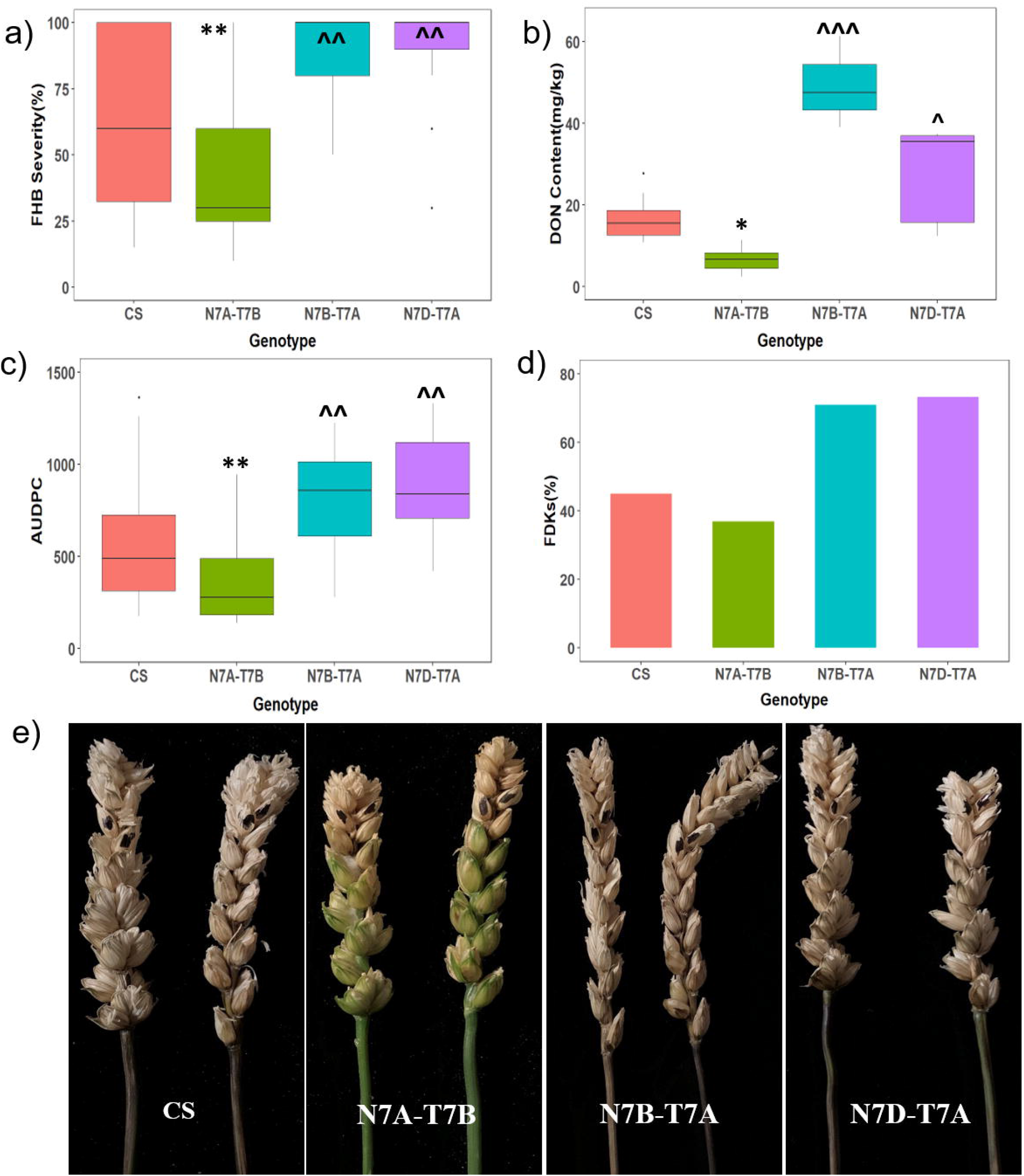
FHB response of Chinese Spring (CS) control and chromosome 7A Nullisomic-tetrasomic lines (X-axis). Y-axis denotes the parameters tested. 1(a): FHB Severity (%), 1(b): DON content(mg/kg); 1(c): AUDPC values; 1(d): FDKs (%); and 1(e): Infected spikes of tested lines (Photographs taken at 28 days after inoculation). * depicts lower significance values than control Chinese Spring at p<0.1, and ** at p<0.01. ^ depicts higher significance values over control Chinese Spring at p<0.1, ^^ at p<0.01, and ^^^ at p<0.001.

For DON content, a one-way ANOVA showed significant genotype effect at p<0.001. N7A-T7B had significantly lower DON at p<0.1 as compared to Chinese spring control. N7B-T7A and N7D-T7A showed significantly higher DON content at p<0.001 and p<0.1, respectively than Chinese spring control (Fig. 1b).

A one-way ANOVA for AUDPC showed significant genotype effect at p<0.001. Compared to control, N7A-T7B were found to have significantly lower AUDPC values whereas N7B-T7A and N7D-T7A had significantly higher values at p<0.01 (Fig. 1c). No variation was observed in the data set for FDK. All Nulli-tetrasomic lines had statistically similar FDKs as that of control at α=0.05 (Fig. 1d).

In the above experiments, since N7A-T7B is missing chromosome 7A and it showed higher level of resistance than control Chinese spring, it indicated the presence of a susceptibility factor on chromosome 7A affecting FHB severity and DON. Furthermore, since N7B-T7A and N7D-T7A plants have four copies of chromosome 7A and thus four doses of putative FHB susceptible factor and these plants, as expected, showed higher level susceptibility to FHB and higher DON content. The results showed that the action of susceptibility gene was affected by chromosome 7A dosage; the deletion of chromosome 7A made the plants resistant, whereas extra copies of 7A made the plants more susceptible to FHB. Therefore, it is justified to call it a FHB susceptibility factor whose deletion leads to resistance.

### FHB susceptibility factor is located on short arm of chromosome 7A (7AS)

Ditelosomic stocks for group-7 chromosomes were evaluated against FHB to map the arm location of the FHB susceptible factor. FHB severity of ditelosomic lines varied from 10 to 100%. At 28 dai, significant genotypic differences at p=2.313e^−10^ with chi-square value of 59.071 were observed. Dt7AL (with short arm of 7A missing) had significantly less FHB severity at p<0.001in contrast to control Chinese spring. All other di-telosomics were found to be statistically similar to control (Fig. 2a; Fig 2e; Supplementary Fig. 1a).

**Figure 2:**
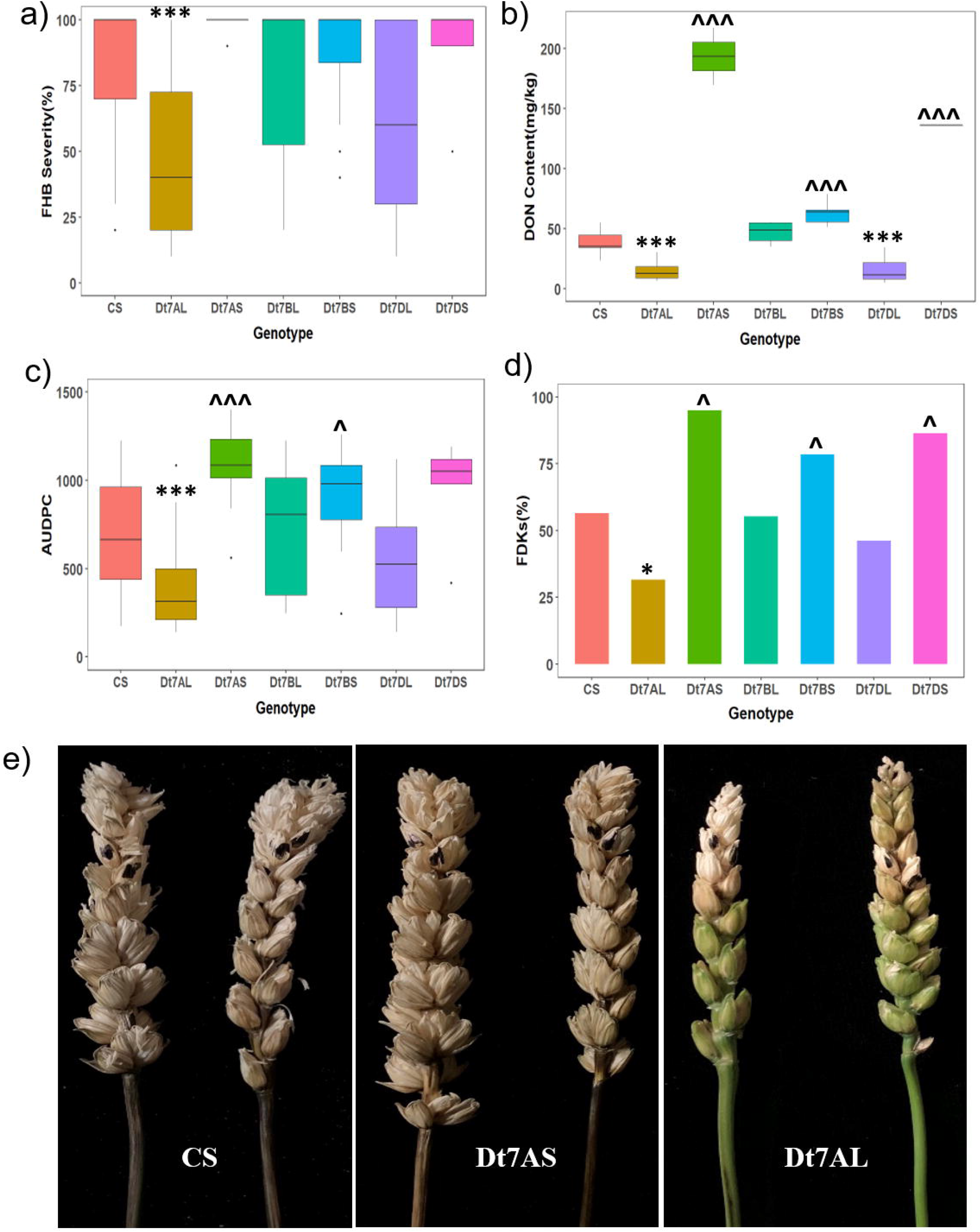
FHB response of the Chinese Spring (CS) control and ditelosomic lines of group-7 chromosomes. X-axis denotes the genotypes and Y-axis denotes the parameters tested. 2(a): FHB Severity (%), 2(b): DON content(mg/kg); 2(c): AUDPC values; 2(d): FDKs(%); and 2(e): Infected spikes of control and ditelosomic lines of 7A (Photographs taken at 28 days after inoculation). * depicts values lower than control Chinese Spring at p<0.05, and *** at p<0.001. ^ depicts higher significance values over control Chinese Spring at p<0.05, and ^^^ at p<0.001.

A one-way ANOVA showed significant genotype effect at p<0.001. Dt7AL and Dt7DL showed significantly lower DON at p<0.001, whereas Dt7AS, Dt7BS and Dt7DS had significantly higher DON content at p<0.001 in comparison to control (Fig. 2b). A one-way ANOVA for AUDPC revealed significant genotypic effect at p<0.001. Dt7AL showed significantly lower AUDPC values at p<0.001, whereas Dt7AS and Dt7BS had significantly higher AUDPC values at p<0.001 and p<0.05 respectively compared to control (Fig. 2c).There was found to be significant variation in the data set for FDK. Dt7AL showed significantly lower FDKs whereas three genotypes (Dt7AS, Dt7BS, Dt7DS) had significantly higher FDKs than control at α=0.05 (Fig. 2d).

Response of ditelosomic lines of group-7 chromosomes revealed the susceptibility factor to be present on short arm of chromosome 7A. Results demonstrated that when the short arm of chromosome 7A is present, plants were susceptible, however, when it was deleted, plants were resistant to FHB.

### The susceptibility factor is conserved in wheat cultivars of diverse origin

In order to test whether this susceptibility factor is conserved across different wheat genotypes or not, six disomic substitution lines derived from five wheat cultivars and one wild tetraploid emmer wheat (chromosome 7A of *T. dicoccoides* substituted chromosome 7A of Chinese spring) were tested for their response to FHB. At 28 dai, a two-way ANOVA showed significant Genotype*Year interaction (p<0.001) in the data set and no significant genotype effect for FHB severity. Therefore, one-way ANOVA was not conducted separately for each year. All substitution lines were found to be statistically indistinguishable from control Chinese spring (Fig. 3a; Fig. 3e).

**Figure 3:**
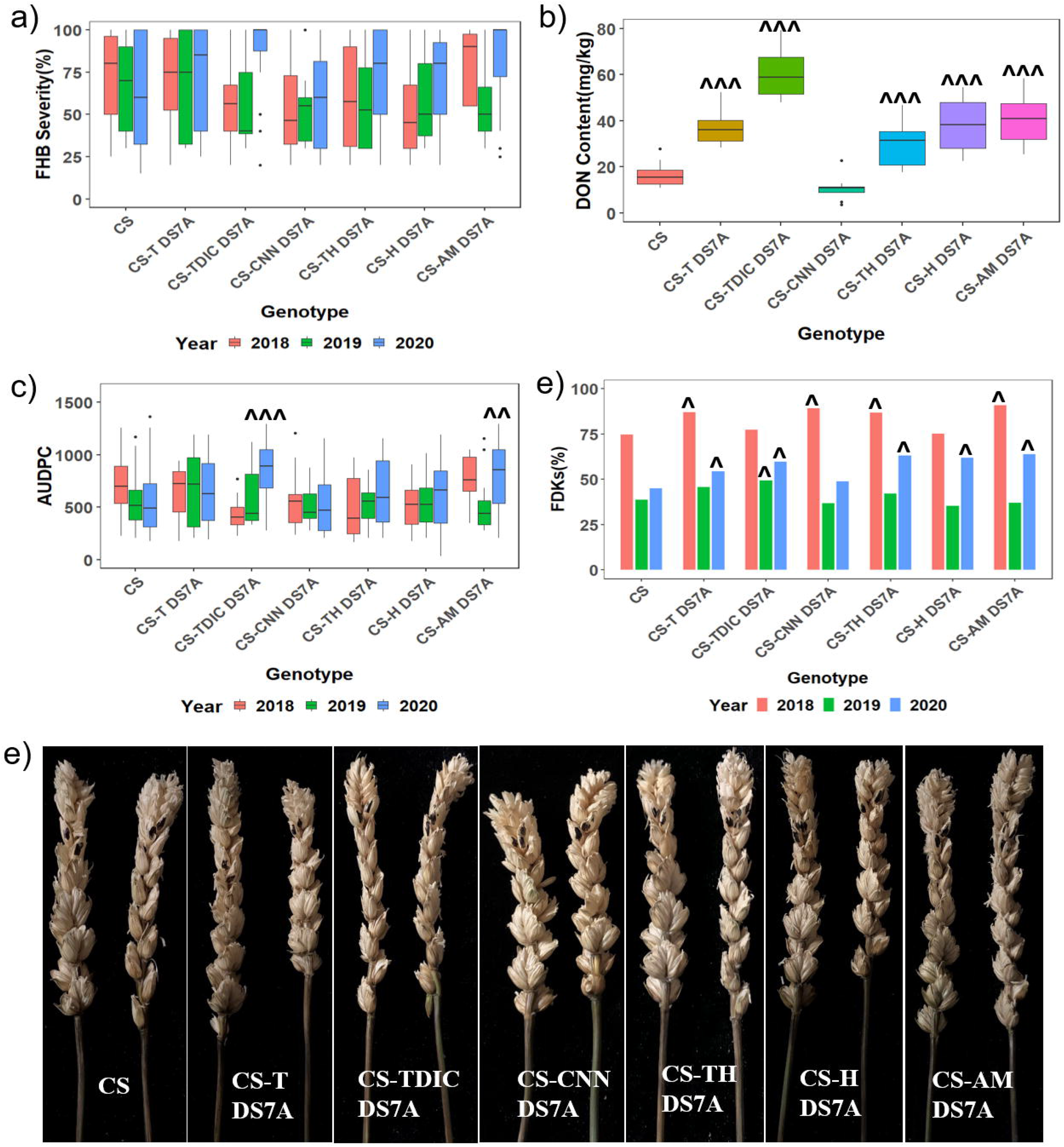
FHB response of the Chinese Spring (CS) control and substitution lines. X-axis denotes the genotypes and Y-axis denotes the parameters tested. 3(a): FHB Severity (%), 3(b): DON content(mg/kg); 3(c): AUDPC values; 3(d): FDKs(%); and 3(e): Infected spikes of tested lines (Photographs taken at 28 days after inoculation). ^ depicts higher significance values over control Chinese Spring at p<0.05, ^^ at p<0.01, and ^^^ at p<0.001.

DON estimation was done for only year 2020. A one-way ANOVA showed significant genotype effect (p<0.001). Five lines showed significantly higher DON content than control. TH had higher DON at p<0.01 and lines (T, H, AMN and TDIC) showed higher DON at p<0.001 (Fig. 3b).

A two-way ANOVA with interaction for all three years of data showed significant genotype effect (p<0.05), year effect (p<0.1) and Genotype*Year interaction (p<0.001) for AUDPC. As G*E effect was found, one-way ANOVA was calculated separately for each year. Data set from years: 2018 (p<0.001) and 2020 (p<0.001) had shown significant genotype effect. For 2018 season, no substitution line was found to be statistically different from control. However, for 2020, two lines (TDIC at p<0.001 and AM at p<0.01) had significantly higher AUDPC values than control. (Fig. 3c).

No substitution line was found to have significantly lower FDKs than control in any of the three testing seasons. Four lines (T, CNN, TH, AM) in 2018, two lines (TDIC) in 2019 and five lines (T, TDIC, TH, H, AM) in 2020 had higher FDKs over control at significance level of α=0.05 (Fig.3d). These results proved that susceptibility factor was conserved across genotypes and was designated as *SF^7AS^FHB*.

### Mapping *SF^7AS^FHB* on Chinese spring reference sequence using deletion lines Molecular characterization of deletion lines

Physical boundaries of the eleven deletion lines of chromosome 7AS were determined using PCR-based molecular markers. Endo & Gill (1996) had sorted these deletion lines on the basis of the fraction length (FL) of chromosome retained using cytogenetic staining techniques as. : 7AS-1 (FL= 0.89), 7AS-9 (FL= 0.89), 7AS-12 (FL= 0.83), 7AS-2 (FL= 0.73), 7AS-5 (FL= 0.59), 7AS-8 (FL= 0.45), 7AS-10 (FL= 0.45), 7AS-11 (FL= 0.33), 7AS-3 (FL= 0.29), 7AS-4 (FL= 0.26), 7AS-6 (FL= 0.21). However, exact Mb position of the deletion lines was not known. To start with, each marker was tested on Chinese spring (positive control), and Nulli 7A (negative control) for confirming the genome specificity. Out of 36 primers, 33 were found to be genome specific for chromosome 7AS. Genome specific markers were then tested on all eleven deletion lines and the results are shown in Table 3. Markers FHB-SF7AS-1, FHB-SF7AS-2 and FHB-SF7AS-3 were found to amplify only control Chinese spring revealing terminal ~30 Mb to be deleted in all the deletion lines. Serially, FHB-SF7AS-4 was the first marker to amplify on del7AS-12 (FL= 0.83), and absent in all other deletion lines, showing del7AS-12 to have retained the maximum segment of chromosome 7AS among the deletion lines. The sizes of deletions for all the lines were deduced in a similar way (Table 3). Deletion line del7AS-10 was found to have small interstitial deletion in addition to its major deletion. Further application of more 7AS-specific PCR markers on del7AS-10 characterized the size of interstitial deletion in it to be ~1Mb in size between 148.4 and 149.2 Mb on reference sequence of 7AS (Supplementary Table 1 and 2). The order of deletion lines was found to be similar with the cytogenetic map of Endo & Gill (1996).

**Table 3:** Marker data on Chinese Spring (Positive control), Nulli 7A (Negative control), and deletion lines. Symbols ‘+’ depicts presence and ‘-’ depicts absence of amplification with the specific primers. Markers are listed in the order of their physical location from the telomere to the centromere and deletion lines are listed based on smallest to largest deletion of terminal 7AS segments.

| Primer/ Genotype | CS | N-7A | 7AS-12 | 7AS-1 | 7AS-9 | 7AS-2 | 7AS-5 | 7AS- 11 | 7AS-10 | 7AS- 8 | 7AS-3 | 7AS-4 | 7AS-6 |
| --- | --- | --- | --- | --- | --- | --- | --- | --- | --- | --- | --- | --- | --- |
| FHB-SF7AS-1 | + | - | - | - | - | - | - | - | - | - | - | - | - |
| FHB-SF7AS-2 | + | - | - | - | - | - | - | - | - | - | - | - | - |
| FHB-SF7AS-3 | + | - | - | - | - | - | - | - | - | - | - | - | - |
| FHB-SF7AS-4 | + | - | + | - | - | - | - | - | - | - | - | - | - |
| FHB-SF7AS-5 | + | - | + | + | + | - | - | - | - | - | - | - | - |
| FHB-SF7AS-6 | + | - | + | + | + | - | - | - | - | - | - | - | - |
| FHB-SF7AS-7 | + | - | + | + | + | - | - | - | - | - | - | - | - |
| FHB-SF7AS-8 | + | - | + | + | + | - | - | - | - | - | - | - | - |
| FHB-SF7AS-9 | + | - | + | + | + | - | - | - | - | - | - | - | - |
| FHB-SF7AS-10 | + | - | + | + | + | + | - | - | - | - | - | - | - |
| FHB-SF7AS-11 | + | - | + | + | + | + | - | - | - | - | - | - | - |
| FHB-SF7AS-12 | + | - | + | + | + | + | - | - | - | - | - | - | - |
| FHB-SF7AS-13 | + | - | + | + | + | + | - | - | - | - | - | - | - |
| FHB-SF7AS-14 | + | - | + | + | + | + | + | - | - | - | - | - | - |
| FHB-SF7AS-15 | + | - | + | + | + | + | + | + | - | - | - | - | - |
| FHB-SF7AS-16 | + | - | + | + | + | + | + | + | - | - | - | - | - |
| FHB-SF7AS-17 | + | - | + | + | + | + | + | + | - | - | - | - | - |
| FHB-SF7AS-18 | + | - | + | + | + | + | + | + | - | - | - | - | - |
| FHB-SF7AS-19 | + | - | + | + | + | + | + | + | - | - | - | - | - |
| FHB-SF7AS-20 | + | - | + | + | + | + | + | + | + | - | - | - | - |
| FHB-SF7AS-21 | + | - | + | + | + | + | + | + | + | - | - | - | - |
| FHB-SF7AS-22 | + | - | + | + | + | + | + | + | - | - | - | - | - |
| FHB-SF7AS-23 | + | - | + | + | + | + | + | + | + | + | - | - | - |
| FHB-SF7AS-24 | + | - | + | + | + | + | + | + | + | + | - | - | - |
| FHB-SF7AS-25 | + | - | + | + | + | + | + | + | + | + | - | - | - |
| FHB-SF7AS-27 | + | - | + | + | + | + | + | + | + | + | + | + | - |
| FHB-SF7AS-28 | + | - | + | + | + | + | + | + | + | + | + | + | - |
| FHB-SF7AS-29 | + | - | + | + | + | + | + | + | + | + | + | + | - |
| FHB-SF7AS-30 | + | - | + | + | + | + | + | + | + | + | + | + | - |
| FHB-SF7AS-32 | + | - | + | + | + | + | + | + | + | + | + | + | + |
| FHB-SF7AS-34 | + | - | + | + | + | + | + | + | + | + | + | + | + |
| FHB-SF7AS-36 | + | - | + | + | + | + | + | + | + | + | + | + | + |

### FHB susceptibility factor is located in the proximal region of 7AS arm

To map *SF^7AS^FHB* to a specific chromosome interval, eleven overlapping deletion lines for short arm of chromosome 7A in Chinese spring background were tested. Control Chinese spring was found to be susceptible in all of the three years. Most of the lines showed similar results for the three years 2018, 2019 and 2020 with some exceptions.

At 28 dai, two-way ANOVA results for FHB severity found significant genotype effect (p<0.001) and Genotype*Year effect (p<0.001) in the data set. One-way ANOVA revealed significant genotype effect for all three seasons (p<0.01 for 2018, p<0.001 for 2019 and 2020). Telomeric, and sub-telomeric deletion lines: del7AS-12, del7AS-9, del7AS-1, del7AS-2, del7AS-5, del7AS-11, del7AS-10 and del7AS-8 were found to have either statistically similar or higher FHB severity as compared to Chinese Spring over all of the three years. Del7AS-10 showed FHB severity statistically similar to Chinese Spring in 2018 and 2019, and lower than control at p<0.01 in 2020. Del7AS-3 showed FHB severity similar to Chinese Spring in year 2018 however, in 2020 it had statistically lower FHB severity than that of Chinese Spring at p<0.001. Del7AS-4 and del7AS-6 showed significantly lower severity in all the years tested (Fig. 4a, Fig. 4e, Supplementary Fig. 1b).

**Figure 4:**
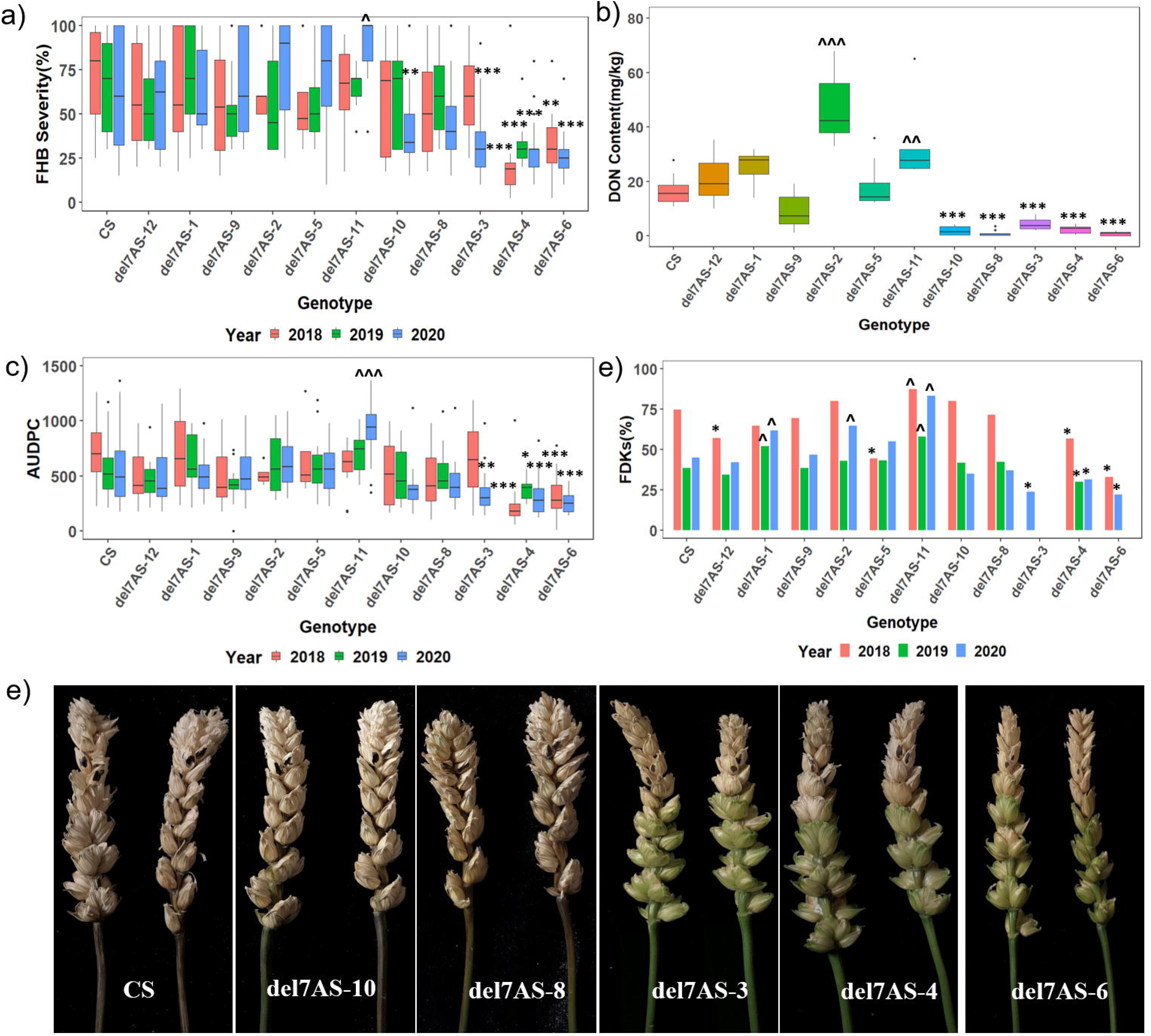
FHB response of Chinese Spring (CS) control and tested deletion lines. X-axis denotes the genotypes and Y-axis denotes the parameters tested. 4(a): FHB Severity (%), 4(b): DON content(mg/kg); 4(c): AUDPC values; 4(d): FDKs (%); and 4(e): Infected spikes of critical deletion lines (Photographs taken at 28 days after inoculation). Lower significance values over control Chinese Spring are depicted with * at p<0.05, ** at p<0.01, and *** at p<0.001. ^ depicts higher significance values over control Chinese Spring at p<0.05, and ^^^ at p<0.001.

DON content measurement was done only for year 2020. One-way ANOVA showed significant genotype effect at p<0.001. Two lines showed significantly higher DON than Chinese Spring (del7AS-11 at p<0.01 and del7AS-2 at p<0.001). Five deletion lines namely del7AS-10, del7AS-8, del7AS-3, del7AS-4, and del7AS-6 were found to have significantly lower DON content at p<0.001 (Fig. 4b). DON content measurement of these critical deletion lines was repeated in another set in 2020 to reconfirm the values.

A two-way ANOVA for all three years data for AUDPC showed significant genotype effect (p<0.001) and Genotype X Year interaction (p<0.001). A separate One-way ANOVA for each year revealed significant genotype effect at p<0.001. In 2018, two genotypes (del7AS-4 and del7AS-6) had significantly lower AUDPC values than Chinese Spring at p<0.001. In 2019, only del7AS-4 showed significantly less AUDPC values at p<0.05. In 2020, three genotypes had significantly lower AUPDC values (del7AS-3 at p<0.01, del7AS-4 and del7AS-6 at p<0.001) whereas del7AS-11 had higher AUPDC values over control Chinese Spring at p<0.001 (Fig. 4c).

In 2018, four genotypes (del7AS-12, del7AS-5, del7AS-4 and del7AS-6) were found to have lower FDKs whereas del7AS-11 had significantly more FDKs in comparison to Chinese Spring at α=0.05. In 2019, one genotype del7AS-4 showed less FDKs at α=0.05 and two genotypes (del7AS-1, del7AS-11) had significantly higher FDKs. In 2020, three genotypes (del7AS-3, del7AS-4, del7AS-6) had significantly lower and three genotypes (del7AS-1, del7AS-2, del7AS-11) had significantly higher FDKs at α=0.05 (Fig. 4d).

These experiments localized the susceptibility factor *SF7^AS^FHB* to the peri-centromeric region of chromosome 7AS. Since peri-centromeric and centromeric deletion lines (del7AS-10, del7AS-8, del7AS-3, del7AS-6, and del7AS-4) were critical in mapping the susceptibility factor and also we lost one year of data (in 2019) for a few of them due to a technical mishap, we tested a set of just these critical lines again in 2020 for their FHB response to robustly locate the susceptibility factor *SF^7AS^FHB*.

### Analysis of critical Deletion lines

At 28 dai, Kruskal-Wallis rank sum test for FHB severity showed significant variation at p= 4.165 e^−06^ with chi-square values of 32.778. Del7AS-6 (p<0.001), del7AS-4 (p<0.05) and del7AS-3 (p<0.01) showed significantly lower severity as compared to control Chinese Spring. Del7AS-8 and del7AS-10 were statistically similar to control (Fig. 5a). For DON content Kruskal-Wallis rank sum test revealed significant variation at p<0.001. Four deletion lines showed lower DON content than control (del7AS-10 and del7AS-4 at p<0.05 whereas del7AS-6 and del7AS-3 at p<0.01). Del7AS-8 had numerically lower, but statistically similar DON content to that of the control Chinese spring (Fig. 5b).

**Figure 5:**
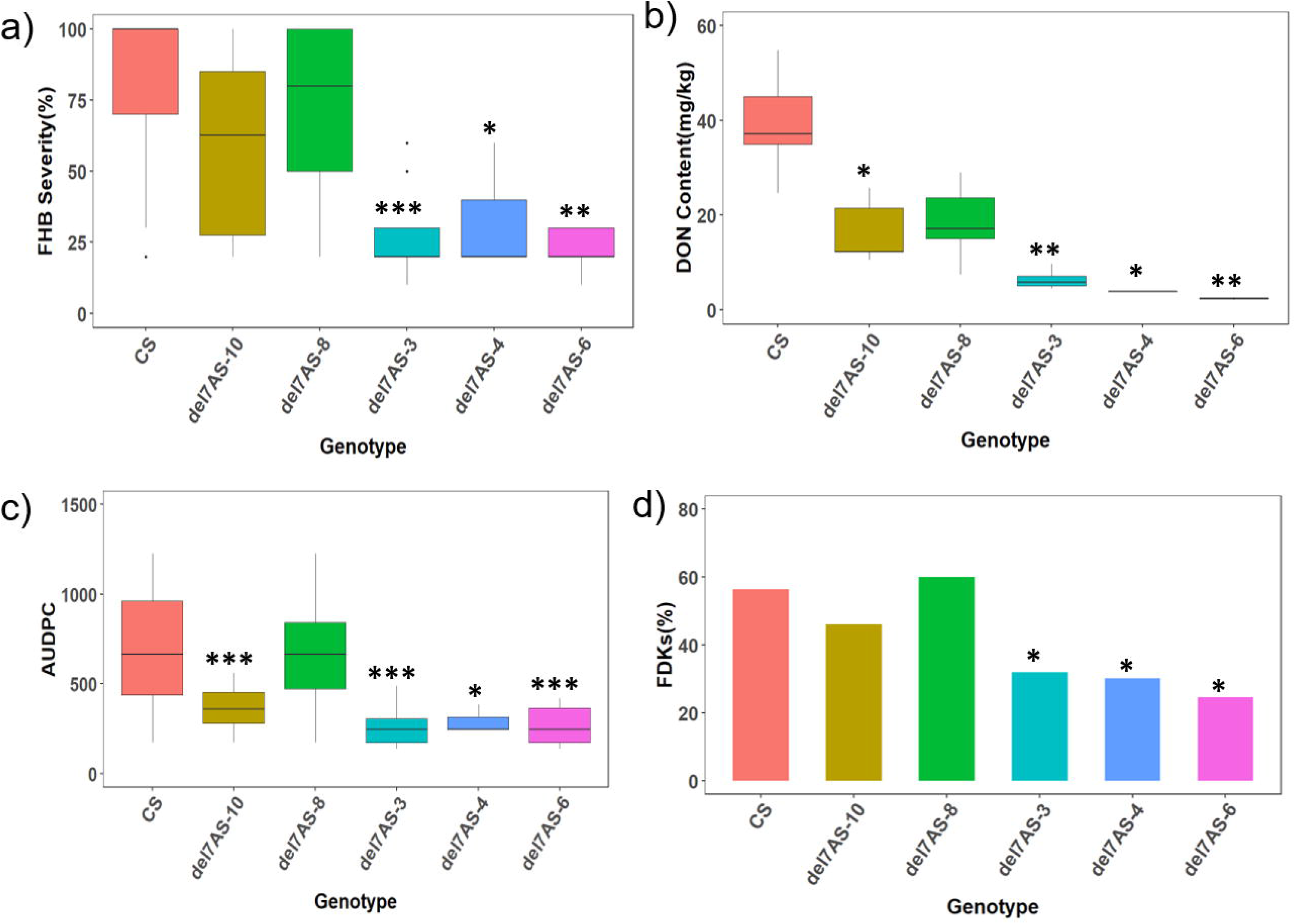
FHB response of Chinese Spring (CS) control and critical deletion lines. X-axis denotes the genotypes and Y-axis denotes the parameters tested. 5(a): FHB Severity (%), 5(b): DON content(mg/kg); 5(c): AUDPC values; 5(d): FDKs(%). * depicts lower significance values over control Chinese Spring at p<0.05, ** at p<0.01, and *** at p<0.001. *7AS*

For AUDPC, a one-way ANOVA showed significant genotype effect at p<0.001. Del7AS-8 was found to be statistically similar to control. Del7AS-10, del7AS-3 and del7AS-6 showed significantly lower AUDPC value at p<0.001, whereas del7AS-4 showed lower disease values at p<0.05 (Fig. 5c). Three genotypes (del7AS-3, del7AS-4, and del7AS-6) showed significantly lower FDKs than control at α=0.05. Del7AS-8 and del7AS-10 were similar to control Chinese spring (Fig. 5d).

### Mapping the susceptibility factor *SF^7AS^FHB*

Del7AS-6, del7AS-4 and del7AS-3 were found to show high level of type-2 (against the spread of the fungal pathogen) and type-4 (towards kernel infection) resistance against FHB, whereas all the other deletion lines were similar to control Chinese spring. Integrated molecular and phenotypic analysis revealed the location of *SF^7AS^FHB* in the ~50 Mb region between 97.5 to 150Mb location on the short arm of chromosome 7A (Figure 6).

**Figure 6:**
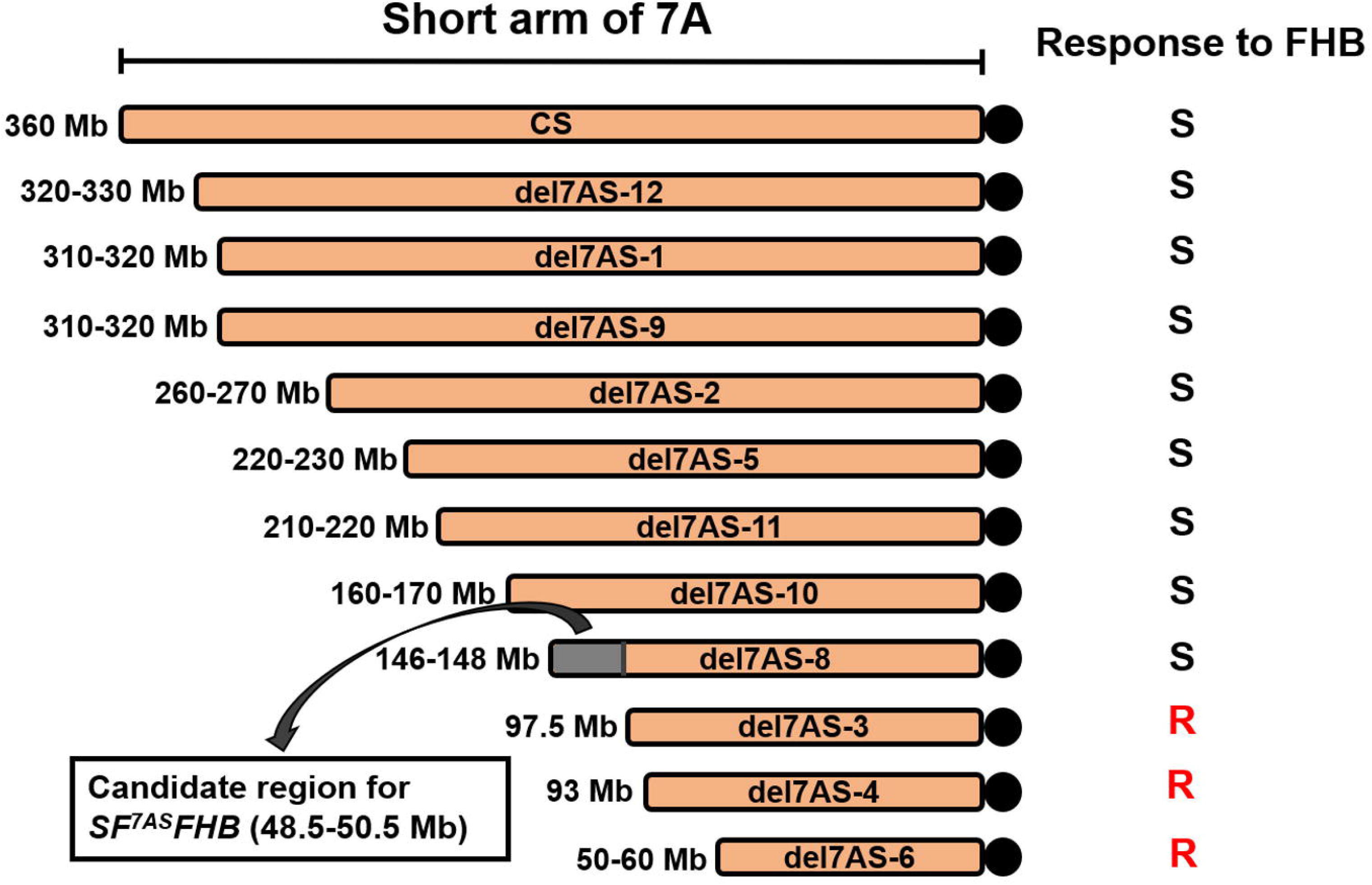
Deletion-bin mapping of the candidate region for susceptibility gene *SF^7AS^FHB*. Deletion lines are depicted in decreasing order of the length of 7A short arm present in them. Response to FHB is shown as S (susceptible) or R (resistant) on right of each of the line. Grey region on 7AS-8 demonstrated the position of *SF^7AS^FHB*.

Some intriguing results were obtained for DON content. Del7AS-3, del7AS-4 and del7AS-6 were found to be resistant for FHB severity, AUDPC and FDKs as well as having low DON content. However, del7AS-10 showed significant susceptibility to FHB severity but significantly lower DON content than control Chinese spring in the two sets (2020 deletion lines and 2020 critical set) analyzed for DON. Critical deletion line del7AS-8 just next to del7AS-3 showed significantly lower DON content than Chinese spring during testing in full set of 2020. When tested again in the subset of critical lines, it was found to be numerically lower, but statistically similar to Chinese spring. It is possible that some additional factor regulating DON content is present in the region of chromosome deleted from del7AS-8 and del7AS-10. However, more experiments are needed to confirm this possibility.

## Discussion

When a pathogen attacks a plant, the result can be a compatible reaction (susceptibility) or an incompatible reaction (resistance). Complex interactions are involved between plant and the pathogen to determine if a plant will be resistant or susceptible. For decades, resistance genes have been known as the major players in imparting resistance to crop plants (Andersen et al., 2018; Kourelis & van der Hoorn, 2018). However, little is known about susceptibility genes facilitating infection by a pathogen in crop plants. Such genes can be good targets for manipulation to make it difficult for the pathogen to survive, and hence making the plants resistant (Engelhardt et al., 2018).

In a study by Ma et al. (2006) a set of 30 ditelosomic lines of Chinese spring was evaluated, and it was found that Dt7AL lacking short arm of chromosome 7A had lowest FHB infection and DON accumulation among all the tested lines. In the present study, we performed systematic deletion bin mapping of this susceptibility factor on chromosome 7AS arm using several critical genetic stocks. We first confirmed the presence of the susceptibility factor on chromosome 7A using nullisomic-tetrasomic lines. This experiment not only showed nullisomic 7A to be resistant to FHB infection, but also revealed a dosage effect of 7A copy number, with N7B-T7A and N7D-T7A to have significantly higher FHB severity and DON content than control Chinese spring. This demonstrated that the identified factor on chromosome 7A works to enhance susceptibility of the plants when present in multiple copies, and hence qualifies as a bonafide susceptibility factor. The factor was named *SF^7AS^ FHB* after confirmation of its presence on the short arm of chromosome 7A.

Using a set of six substitution lines from different varieties/ species of wheat, we found that the *SF^7AS^ FHB* gene is conserved across not only different varieties, but also in tetraploid species. All tested substitution lines were found to be statistically similar or showing higher disease susceptibility in comparison to Chinese Spring for all the four tested parameters. This indicates that the susceptibility factor is retained by most of the varieties/ species of wheat. Hence, developing a bi-parental genetic mapping population for mapping this trait would not be feasible.

We used 11 overlapping deletion lines of 7AS chromosome arm to map the location of the susceptibility gene *FHB-SF^7AS^*. In the first step, we precisely characterized all the deletion sizes using genome specific molecular markers based on the Wheat Reference Genome Sequence v 1.0 (IWGSC et al. 2018). Each deletion line was ordered according to their deletion size, and the order was found to be in agreement with that reported by Endo & Gill (1996).

We tested response of deletion lines to FHB four times in the greenhouse and did estimation of four parameters. For FHB Severity, AUDPC, and FDKs, three pericentromeric lines: 7AS-6, 7AS-4 and 7AS-3 were found to be consistently resistant whereas all other lines were found to be susceptible. Comparative molecular and phenotypic analysis revealed the *SF^7AS^FHB* to be present between del7AS-3 (resistant) and del7AS-8 (susceptible) in a peri-centromeric region of 48.5-50.5 Mb. This indicates that deletion lines (del7AS-12, del7AS-1, del7AS-9, del7AS-2, del7AS-5, del7AS-11, del7AS-10 and del7AS-8) harbor the susceptibility gene *SF^7AS^FHB* and loss of the chromosome segment carrying the susceptibility gene (*SF^7AS^FHB*) makes the centromeric lines (del7AS-3, del7AS-4, and del7AS-6) resistant. These three parameters reinforce our conclusion of the candidate region and show *SF^7AS^FHB* to be affecting Type-2 (resistance to spread of the pathogen) and Type-4 (reduced kernel infection) resistance.

We also estimated DON content of nulli-tetrasomic, ditelosomic lines, substitution and deletion lines. Except for nulli-tetrasomic lines, we found differences in results for this parameter as compared to the results of the other three tested parameters. For di-telosomic lines, we found that in addition to Dt 7AL, Dt 7DL also had significantly lower DON content than control Chinese spring, however, Dt 7DL was statistically similar to Chinese spring for FHB severity, AUDPC and FDKs. This might be due to 7DS harboring a susceptibility gene imparting resistance against DON (Type-3 resistance). All six substitution lines tested were found to have DON content similar to control Chinese spring. Deletion lines del7AS-6, del7AS-4, del7AS-3, del7AS-8, and del7AS-10 had less DON content, whereas all other deletion lines had high DON content values. It is possible that an additional gene regulating DON content is located in the region between del7AS-10 and del7AS-11. Possibly, the 1Mb interstitial deletion in del7AS-10 (Supplementary Table 1), which is missing from other resistant lines as well as del7AS-8 harbors such a factor regulating DON content. However, more experiments are needed for validation of this proposal. Hales et al. (2020) also reported presence of two different susceptibility genes on 4DS, one providing resistance against FHB severity and the other against DON content.

It is worth mentioning here that deletion lines did not show any major difference in phenotypes from control Chinese Spring over all the three seasons. Deletion line 7AS-11 was found to be susceptible, but also had some other differences in morphology in terms of late flowering habit and having compact and light green spikes. It was susceptible and hence, does contain susceptibility gene. However, 7AS-11 was reported to have one more deletion in short arm of chromosome 2B (Endo and Gill, 1996), which appears to be the reason of abnormal morphology of this deletion line. The mapped region of 48.5-50.5 Mb interval is still a big interval and needs to be refined further. To do that, we plan to use Gamma-irradiated Radiation Hybrid Panel of Chinese spring to detect smaller deletions in the candidate region and delineate the region further (Tiwari et al., 2016).

Analysis of diverse substitution lines showed *FHB-SF^7AS^* to be conserved across varieties and even distinct species. It is known that genetic recombination events in wheat are limited to sub-telomeric regions. For example, in chromosome 3B of wheat, 82% of the cross-over events are restricted to the distal ends of the chromosome, representing only 19% of the whole chromosome length (Saintenac et al., 2009; Darrier et al., 2017). *FHB-SF^7AS^* is located in the peri-centromeric region of 7AS and hence, its conserved nature is according to the expectations.

Pathogens are known to hijack/utilize plant machinery for their own benefit and they target susceptibility genes/ resistance suppressors for this purpose. As mentioned earlier, these genes can help a pathogen either in its establishment or has a role in regulating plant defenses in favor of pathogen or supports pathogen living on the host (Lapin & Van den Ackerveken, 2013). Nitrogen fixation is an example of a susceptibility gene. Though, it has symbiotic relationship between plant and bacteria, but bacteria are utilizing plant genes for their survival. Plants release flavonoids into soil which acts a chemoattractant for attracting Rhizobium (Hirsch, 1992). The *mlo* locus (mildew resistance locus O), the earliest and widely studied susceptibility gene, is used by powdery mildew fungus to penetrate into epidermal cells of the host (Consonni et al., 2006; Miklis et al., 2007). *ROP GTPase RACB* of Barley assists in the expansion of haustoria of powdery mildew fungus in leaf epidermal cells (Hoefle et al., 2011; Scheler et al., 2016). *PMR6*, a pectate lyase degrades pectin in the cell wall and supports powdery mildew fungus growth on Arabidopsis leaves (Vogel et al., 2002). Faris et al. (2010) reported *Tsn1* gene in wheat to govern effector-triggered susceptibility to necrotrophic fungal pathogens *Stagonospora nodorum* and *Pyrenophora tritici-repentis.* Fine mapping and identification of *SF^7AS^FHB* gene will allow better understanding of its role in the plant and in the interaction with *F. graminearum* and further its utilization in wheat varieties.

Only a few attempts at mapping susceptibility genes in wheat against *F.graminearum* have been reported till date. Wheat *RPL3* gene family assists *F. graminearum* in production of virulence factor DON and mutating *RPL3* imparts resistance against FHB (Lucyshyn et al., 2007). *Ethylene Insensitive 2* (*EIN2*), a gene in wheat involved in Ethylene signaling was found to support *F. graminearum* for its growth. *ein2* mutants had reduced disease severity and DON content, suggesting possible manipulation of this gene and ethylene signaling pathway by the fungus (Chen et al., 2009). Silencing the *TaLpx-1* gene, which encodes 9-lipoxygenase, rendered wheat resistant to *F. graminearum* (Nalam et al., 2015). Garvin et al. (2015) reported a susceptibility gene located on chromosome 3DL of wheat. However, fine mapping of that region has not been reported till date. Hales et al. (2020) identified a susceptibility gene for FHB on chromosome 4DS and mapped it to a 31.7 Mbp interval. In the present study, we identified a new susceptibility factor (*SF^7AS^FHB*) on chromosome 7A short arm. We were able to map the region of the susceptibility gene *SF^7AS^FHB* to 48.5-50.5 Mbp.

Analysis of all the parameters revealed that removal of susceptibility gene region provides 50-60 % resistance against FHB. No obvious phenological or morphological penalty of deletion of SF^7AS^FHB was observed in the resistant deletion lines. The effect of *SF^7AS^FHB* region was found to be conserved across the genotypes, giving broader and durable prospects to resistance conferred by manipulation of the 7AS susceptibility gene.

## Acknowledgements

Authors are thankful to US Wheat Barley and Scab Initiative (Award# 59-0206-0-177, 59-0200-6-018), National Science Foundation (Award# 1943155) and USDA NIFA (Award# 2020-67013-32558 and 2020-67013-31460) for financial support.

## References

Andersen, E. J., Ali, S., Byamukama, E., Yen, Y., & Nepal, M. P. (2018). Disease Resistance Mechanisms in Plants. Genes, 9(7). https://doi.org/10.3390/genes9070339

Bai, G., & Shaner, G. (2004). Management and resistance in wheat and barley to fusarium head blight. Annual Review of Phytopathology, 42(1), 135–161. https://doi.org/10.1146/annurev.phyto.42.040803.140340

Bai, G., & Shaner, G. E. (1994). Scab of wheat: Prospects for control. https://doi.org/10.1094/PD-78-0760

Bai, G.-H., Desjardins, A. E., & Plattner, R. D. (2002). Deoxynivalenol-nonproducing Fusarium graminearum Causes Initial Infection, but does not Cause DiseaseSpread in Wheat Spikes. Mycopathologia, 153(2), 91–98. https://doi.org/10.1023/A:1014419323550

Buerstmayr, M., Steiner, B., & Buerstmayr, H. (n.d.). Breeding for Fusarium head blight resistance in wheat—Progress and challenges. Plant Breeding, *n/a*(n/a). https://doi.org/10.1111/pbr.12797

Büschges, R., Hollricher, K., Panstruga, R., Simons, G., Wolter, M., Frijters, A., van Daelen, R., van der Lee, T., Diergaarde, P., Groenendijk, J., Töpsch, S., Vos, P., Salamini, F., & Schulze-Lefert, P. (1997). The Barley Mlo Gene: A Novel Control Element of Plant Pathogen Resistance. Cell, 88(5), 695–705. https://doi.org/10.1016/S0092-8674(00)81912-1

Cainong, J. C., Bockus, W. W., Feng, Y., Chen, P., Qi, L., Sehgal, S. K., Danilova, T. V., Koo, D.-H., Friebe, B., & Gill, B. S. (2015). Chromosome engineering, mapping, and transferring of resistance to Fusarium head blight disease from Elymus tsukushiensis into wheat. Theoretical and Applied Genetics, 128(6), 1019–1027. https://doi.org/10.1007/s00122-015-2485-1

Chen, X., Steed, A., Travella, S., Keller, B., & Nicholson, P. (2009). Fusarium graminearum exploits ethylene signalling to colonize dicotyledonous and monocotyledonous plants. New Phytologist, 182(4), 975–983. https://doi.org/10.1111/j.1469-8137.2009.02821.x

Chen, Y., Kistler, H. C., & Ma, Z. (2019). Fusarium graminearum Trichothecene Mycotoxins: Biosynthesis, Regulation, and Management. Annual Review of Phytopathology, 57(1), 15–39. https://doi.org/10.1146/annurev-phyto-082718-100318

Consonni, C., Humphry, M. E., Hartmann, H. A., Livaja, M., Durner, J., Westphal, L., Vogel, J., Lipka, V., Kemmerling, B., Schulze-Lefert, P., Somerville, S. C., & Panstruga, R. (2006). Conserved requirement for a plant host cell protein in powdery mildew pathogenesis. Nature Genetics, 38(6), 716–720. https://doi.org/10.1038/ng1806

Consortium (IWGSC), T. I. W. G. S., Appels, R., Eversole, K., Stein, N., Feuillet, C., Keller, B., Rogers, J., Pozniak, C. J., Choulet, F., Distelfeld, A., Poland, J., Ronen, G., Sharpe, A. G., Barad, O., Baruch, K., Keeble-Gagnère, G., Mascher, M., Ben-Zvi, G., Josselin, A.-A., … Wang, L. (2018a). Shifting the limits in wheat research and breeding using a fully annotated reference genome. Science, 361(6403). https://doi.org/10.1126/science.aar7191

Consortium (IWGSC), T. I. W. G. S., Appels, R., Eversole, K., Stein, N., Feuillet, C., Keller, B., Rogers, J., Pozniak, C. J., Choulet, F., Distelfeld, A., Poland, J., Ronen, G., Sharpe, A. G., Barad, O., Baruch, K., Keeble-Gagnère, G., Mascher, M., Ben-Zvi, G., Josselin, A.-A., … Wang, L. (2018b). Shifting the limits in wheat research and breeding using a fully annotated reference genome. Science, 361(6403). https://doi.org/10.1126/science.aar7191

Cuthbert, P. A., Somers, D. J., & Brulé-Babel, A. (2007). Mapping of Fhb2 on chromosome 6BS: A gene controlling Fusarium head blight field resistance in bread wheat (Triticum aestivum L.). Theoretical and Applied Genetics, 114(3), 429–437. https://doi.org/10.1007/s00122-006-0439-3

Cuthbert, P. A., Somers, D. J., Thomas, J., Cloutier, S., & Brulé-Babel, A. (2006). Fine mapping Fhb1, a major gene controlling fusarium head blight resistance in bread wheat (Triticum aestivum L.). Theoretical and Applied Genetics, 112(8), 1465. https://doi.org/10.1007/s00122-006-0249-7

Cutler, H. G. (1988). Trichothecenes and Their Role in the Expression of Plant Disease. In Biotechnology for Crop Protection (Vol. 379, pp. 50–72). American Chemical Society. https://doi.org/10.1021/bk-1988-0379.ch004

Darrier, B., Rimbert, H., Balfourier, F., Pingault, L., Josselin, A.-A., Servin, B., Navarro, J., Choulet, F., Paux, E., & Sourdille, P. (2017). High-Resolution Mapping of Crossover Events in the Hexaploid Wheat Genome Suggests a Universal Recombination Mechanism. Genetics, 206(3), 1373–1388. https://doi.org/10.1534/genetics.116.196014

Desjardins, A. E. (usda/Ars, Proctor, R. H., McCormick, S. P., & Hohn, T. M. (1997). Reduced virulence of trichothecene antibiotic-nonproducing mutants of Gibberella zeae in wheat field tests. Proceedings of a Workshop, El Batan, Mex. (Mexico), 13-17 Oct 1996. https://agris.fao.org/agris-search/search.do?recordID=QY1998000294

Eckardt, N. A. (2002). Plant Disease Susceptibility Genes? The Plant Cell, 14(9), 1983–1986. https://doi.org/10.1105/tpc.140910

Endo, T. R., & Gill, B. S. (1996). The Deletion Stocks of Common Wheat. Journal of Heredity, 87(4), 295–307. https://doi.org/10.1093/oxfordjournals.jhered.a023003

Endo, T.R. (1986). Complete Identification of common wheat chromosomes by means of the C-banding technique. Japanese Journal of Genetics, 61, 89–93.

Endo, T.R. (2015). New Aneuploids of Common Wheat. In Y. Ogihara, S. Takumi, & H. Handa (Eds.), Advances in Wheat Genetics: From Genome to Field (pp. 73–81). Springer Japan. https://doi.org/10.1007/978-4-431-55675-6_8

Engelhardt, S., Stam, R., & Hückelhoven, R. (2018). Good Riddance? Breaking Disease Susceptibility in the Era of New Breeding Technologies. Agronomy, 8(7), 114. https://doi.org/10.3390/agronomy8070114

Gao, L., Diarso, M., Zhang, A., Zhang, H., Dong, Y., Liu, L., Lv, Z., & Liu, B. (2016). Heritable alteration of DNA methylation induced by whole-chromosome aneuploidy in wheat. The New Phytologist, 209(1), 364–375. https://doi.org/10.1111/nph.13595

Goswami, R. S., & Kistler, H. C. (2004). Heading for disaster: Fusarium graminearum on cereal crops. Molecular Plant Pathology, 5(6), 515–525. https://doi.org/10.1111/j.1364-3703.2004.00252.x

Guo, J., Zhang, X., Hou, Y., Cai, J., Shen, X., Zhou, T., Xu, H., Ohm, H. W., Wang, H., Li, A., Han, F., Wang, H., & Kong, L. (2015). High-density mapping of the major FHB resistance gene Fhb7 derived from Thinopyrum ponticum and its pyramiding with Fhb1 by marker-assisted selection. Theoretical and Applied Genetics, 128(11), 2301–2316. https://doi.org/10.1007/s00122-015-2586-x

Gupta, P. K., & Vasistha, N. K. (2018). Wheat cytogenetics and cytogenomics: The present status. The Nucleus, 61(3), 195–212. https://doi.org/10.1007/s13237-018-0243-x

Hales, B., Steed, A., Giovannelli, V., Burt, C., Lemmens, M., Molnár-Láng, M., & Nicholson, P. (n.d.). Type II Fusarium head blight susceptibility conferred by a region on wheat chromosome 4D. Journal of Experimental Botany. https://doi.org/10.1093/jxb/eraa226

Han, X., & Kahmann, R. (2019). Manipulation of Phytohormone Pathways by Effectors of Filamentous Plant Pathogens. Frontiers in Plant Science, 10. https://doi.org/10.3389/fpls.2019.00822

Hirsch, A. M. (1992). Tansley Review No. 40. Developmental Biology of Legume Nodulation. The New Phytologist, 122(2), 211–237. JSTOR.

Hoefle, C., Huesmann, C., Schultheiss, H., Börnke, F., Hensel, G., Kumlehn, J., & Hückelhoven, R. (2011). A Barley ROP GTPase ACTIVATING PROTEIN Associates with Microtubules and Regulates Entry of the Barley Powdery Mildew Fungus into Leaf Epidermal Cells. The Plant Cell, 23(6), 2422–2439. https://doi.org/10.1105/tpc.110.082131

Jansen, C., Wettstein, D. von, Schäfer, W., Kogel, K.-H., Felk, A., & Maier, F. J. (2005). Infection patterns in barley and wheat spikes inoculated with wild-type and trichodiene synthase gene disrupted Fusarium graminearum. Proceedings of the National Academy of Sciences, 102(46), 16892–16897. https://doi.org/10.1073/pnas.0508467102

Kazan, K., & Lyons, R. (2014). Intervention of Phytohormone Pathways by Pathogen Effectors. The Plant Cell, 26(6), 2285–2309. https://doi.org/10.1105/tpc.114.125419

Kourelis, J., & van der Hoorn, R. A. L. (2018). Defended to the Nines: 25 Years of Resistance Gene Cloning Identifies Nine Mechanisms for R Protein Function[OPEN]. The Plant Cell, 30(2), 285–299. https://doi.org/10.1105/tpc.17.00579

Lapin, D., & Van den Ackerveken, G. (2013). Susceptibility to plant disease: More than a failure of host immunity. Trends in Plant Science, 18(10), 546–554. https://doi.org/10.1016/j.tplants.2013.05.005

Lucyshyn, D., Busch, B. L., Abolmaali, S., Steiner, B., Chandler, E., Sanjarian, F., Mousavi, A., Nicholson, P., Buerstmayr, H., & Adam, G. (2007). Cloning and characterization of the ribosomal protein L3 (RPL3) gene family from Triticum aestivum. Molecular Genetics and Genomics, 277(5), 507–517. https://doi.org/10.1007/s00438-006-0201-1

Ma, H.-X., Bai, G.-H., Gill, B. S., & Hart, L. P. (2006). Deletion of a Chromosome Arm Altered Wheat Resistance to Fusarium Head Blight and Deoxynivalenol Accumulation in Chinese Spring. Plant Disease, 90(12), 1545–1549. https://doi.org/10.1094/PD-90-1545

McMullen, M., Jones, R., & Gallenberg, D. (1997). Scab of Wheat and Barley: A Re-emerging Disease of Devastating Impact. Plant Disease, 81(12), 1340–1348. https://doi.org/10.1094/PDIS.1997.81.12.1340

Mesterházy, Á., Bartók, T., Mirocha, C. G., & Komoróczy, R. (1999). Nature of wheat resistance to Fusarium head blight and the role of deoxynivalenol for breeding. Plant Breeding, 118(2), 97–110. https://doi.org/10.1046/j.1439-0523.1999.118002097.x

Miklis, M., Consonni, C., Bhat, R. A., Lipka, V., Schulze-Lefert, P., & Panstruga, R. (2007). Barley MLO Modulates Actin-Dependent and Actin-Independent Antifungal Defense Pathways at the Cell Periphery. Plant Physiology, 144(2), 1132–1143. https://doi.org/10.1104/pp.107.098897

Mirocha, C. J., Kolaczkowski, E., Xie, W., Yu, H., & Jelen, H. (1998). Analysis of Deoxynivalenol and Its Derivatives (Batch and Single Kernel) Using Gas Chromatography/Mass Spectrometry. Journal of Agricultural and Food Chemistry, 46(4), 1414–1418. https://doi.org/10.1021/jf970857o

Nalam, V. J., Alam, S., Keereetaweep, J., Venables, B., Burdan, D., Lee, H., Trick, H. N., Sarowar, S., Makandar, R., & Shah, J. (2015). Facilitation of Fusarium graminearum Infection by 9-Lipoxygenases in Arabidopsis and Wheat. Molecular Plant-Microbe Interactions®, 28(10), 1142–1152. https://doi.org/10.1094/MPMI-04-15-0096-R

Parry, D. W., Jenkinson, P., & McLEOD, L. (1995a). Fusarium ear blight (scab) in small grain cereals—A review. Plant Pathology, 44(2), 207–238. https://doi.org/10.1111/j.1365-3059.1995.tb02773.x

Parry, D. W., Jenkinson, P., & McLEOD, L. (1995b). Fusarium ear blight (scab) in small grain cereals?a review. Plant Pathology, 44(2), 207–238. https://doi.org/10.1111/j.1365-3059.1995.tb02773.x

Pavan, S., Jacobsen, E., Visser, R. G. F., & Bai, Y. (2009). Loss of susceptibility as a novel breeding strategy for durable and broad-spectrum resistance. Molecular Breeding, 25(1), 1. https://doi.org/10.1007/s11032-009-9323-6

Qi, L. L., Pumphrey, M. O., Friebe, B., Chen, P. D., & Gill, B. S. (2008). Molecular cytogenetic characterization of alien introgressions with gene Fhb3 for resistance to Fusarium head blight disease of wheat. Theoretical and Applied Genetics, 117(7), 1155–1166. https://doi.org/10.1007/s00122-008-0853-9

Raupp, J. W. (1995). Suggested guidelines for the nomenclature and abbreviation of the genetic stocks of wheat, Triticum aestivum L.em Thell., and its relatives. Wheat Information Service, 81, 50–55.

Rawat, N., Pumphrey, M. O., Liu, S., Zhang, X., Tiwari, V. K., Ando, K., Trick, H. N., Bockus, W. W., Akhunov, E., Anderson, J. A., & Gill, B. S. (2016). Wheat Fhb1 encodes a chimeric lectin with agglutinin domains and a pore-forming toxin-like domain conferring resistance to Fusarium head blight. Nature Genetics, 48(12), 1576–1580. https://doi.org/10.1038/ng.3706

Rocha, O., Ansari, K., & Doohan, F. M. (2005). Effects of trichothecene mycotoxins on eukaryotic cells: A review. Food Additives and Contaminants, 22(4), 369–378. https://doi.org/10.1080/02652030500058403

Saintenac, C., Falque, M., Martin, O. C., Paux, E., Feuillet, C., & Sourdille, P. (2009). Detailed Recombination Studies Along Chromosome 3B Provide New Insights on Crossover Distribution in Wheat (Triticum aestivum L.). Genetics, 181(2), 393–403. https://doi.org/10.1534/genetics.108.097469

Salgado, J. D., Madden, L. V., & Paul, P. A. (2014). Efficacy and Economics of Integrating In-Field and Harvesting Strategies to Manage Fusarium Head Blight of Wheat. Plant Disease, 98(10), 1407–1421. https://doi.org/10.1094/PDIS-01-14-0093-RE

Scheler, B., Schnepf, V., Galgenmüller, C., Ranf, S., & Hückelhoven, R. (2016). Barley disease susceptibility factor RACB acts in epidermal cell polarity and positioning of the nucleus. Journal of Experimental Botany, 67(11), 3263–3275. https://doi.org/10.1093/jxb/erw141

Sears, E. R. (1966). Nullisomic-Tetrasomic Combinations in Hexaploid Wheat. In R. Riley & K. R. Lewis (Eds.), Chromosome Manipulations and Plant Genetics: The contributions to a symposium held during the Tenth International Botanical Congress Edinburgh 1964 (pp. 29–45). Springer US. https://doi.org/10.1007/978-1-4899-6561-5_4

Sears, Ernest Robert. (1954). The aneuploids of common wheat. University of Missouri, College of Agriculture, Agricultural Experiment Station.

Sears: The aneuploids of common wheat—Google Scholar. (n.d.). Retrieved July 25, 2020, from https://scholar-google-com.proxy-um.researchport.umd.edu/scholar_lookup?title=The%20aneuploids%20of%20common%20wheat&author=ER.%20Sears&journal=Missouri%20Agr%20Expt%20Sta%20Res%20Bull&volume=572&pages=1-59&publication_year=1954

Simko, I., & Piepho, H.-P. (2011). The Area Under the Disease Progress Stairs: Calculation, Advantage, and Application. Phytopathology®, 102(4), 381–389. https://doi.org/10.1094/PHYTO-07-11-0216

Snijders, C. H. A. (1990). Fusarium head blight and mycotoxin contamination of wheat, a review. Netherlands Journal of Plant Pathology, 96(4), 187–198. https://doi.org/10.1007/BF01974256

Song, Q. J., Shi, J. R., Singh, S., Fickus, E. W., Costa, J. M., Lewis, J., Gill, B. S., Ward, R., & Cregan, P. B. (2005). Development and mapping of microsatellite (SSR) markers in wheat. Theoretical and Applied Genetics, 110(3), 550–560. https://doi.org/10.1007/s00122-004-1871-x

Tiwari, V. K., Heesacker, A., RieraLLizarazu, O., Gunn, H., Wang, S., Wang, Y., Gu, Y. Q., Paux, E., Koo, D.-H., Kumar, A., Luo, M.-C., Lazo, G., Zemetra, R., Akhunov, E., Friebe, B., Poland, J., Gill, B. S., Kianian, S., & Leonard, J. M. (2016). A whole-genome, radiation hybrid mapping resource of hexaploid wheat. The Plant Journal, 86(2), 195–207. https://doi.org/10.1111/tpj.13153

van Schie, C. C. N., & Takken, F. L. W. (2014). Susceptibility Genes 101: How to Be a Good Host. Annual Review of Phytopathology, 52(1), 551–581. https://doi.org/10.1146/annurev-phyto-102313-045854

Vogel, J. P., Raab, T. K., Schiff, C., & Somerville, S. C. (2002). PMR6, a pectate lyase-like gene required for powdery mildew susceptibility in Arabidopsis. The Plant Cell, 14(9), 2095–2106. https://doi.org/10.1105/tpc.003509

Waldron, B. L., MorenoLSevilla, B., Anderson, J. A., Stack, R. W., & Frohberg, R. C. (1999). RFLP Mapping of QTL for Fusarium Head Blight Resistance in Wheat. Crop Science, 39(3), cropsci1999.0011183X003900030032x. https://doi.org/10.2135/cropsci1999.0011183X003900030032x

Wang, H., Sun, S., Ge, W., Zhao, L., Hou, B., Wang, K., Lyu, Z., Chen, L., Xu, S., Guo, J., Li, M., Su, P., Li, X., Wang, G., Bo, C., Fang, X., Zhuang, W., Cheng, X., Wu, J., … Kong, L. (2020). Horizontal gene transfer of Fhb7 from fungus underlies Fusarium head blight resistance in wheat. Science. https://doi.org/10.1126/science.aba5435

Wang, Y., Tiwari, V. K., Rawat, N., Gill, B. S., Huo, N., You, F. M., Coleman-Derr, D., & Gu, Y. Q. (2016). GSP: A web-based platform for designing genome-specific primers in polyploids. Bioinformatics, 32(15), 2382–2383. https://doi.org/10.1093/bioinformatics/btw134

Wegulo, S. N., Bockus, W. W., Nopsa, J. H., De Wolf, E. D., Eskridge, K. M., Peiris, K. H. S., & Dowell, F. E. (2010). Effects of Integrating Cultivar Resistance and Fungicide Application on Fusarium Head Blight and Deoxynivalenol in Winter Wheat. Plant Disease, 95(5), 554–560. https://doi.org/10.1094/PDIS-07-10-0495

Werner JE, Endo TR, and Gill BS, 1992. Toward a cytogenetically based physical map of the wheat genome. Proceedings of the National Academy of Sciences, USA, 8,11307–11311.

Wilson, W., Dahl, B., & Nganje, W. (2018). Economic costs of Fusarium Head Blight, scab and deoxynivalenol. World Mycotoxin Journal, 11(2), 291–302. https://doi.org/10.3920/WMJ2017.2204

Xue, S., Li, G., Jia, H., Xu, F., Lin, F., Tang, M., Wang, Y., An, X., Xu, H., Zhang, L., Kong, Z., & Ma, Z. (2010). Fine mapping Fhb4, a major QTL conditioning resistance to Fusarium infection in bread wheat (Triticum aestivum L.). Theoretical and Applied Genetics, 121(1), 147–156. https://doi.org/10.1007/s00122-010-1298-5

Xue, S., Xu, F., Tang, M., Zhou, Y., Li, G., An, X., Lin, F., Xu, H., Jia, H., Zhang, L., Kong, Z., & Ma, Z. (2011). Precise mapping Fhb5, a major QTL conditioning resistance to Fusarium infection in bread wheat (Triticum aestivum L.). Theoretical and Applied Genetics, 123(6), 1055–1063. https://doi.org/10.1007/s00122-011-1647-z

